# Retinoic acid signaling alters the balance of epidermal stem cell populations in the skin

**DOI:** 10.1101/2025.04.10.648290

**Authors:** Thisakorn Dumrongphuttidecha, Mizuho Ishikawa, Nina Cabezas-Wallscheid, Aiko Sada

## Abstract

All-trans retinoic acid (ATRA), the active form of retinoids, is a potent anti-aging and anti-inflammatory agent with pleiotropic effects on various skin conditions. In the interfollicular epidermis, tissue turnover is maintained by heterogeneous epidermal stem cell populations located at the basal layer. Mouse tail skin contains slow- and fast-cycling epidermal stem cell populations. The reduction of fast-cycling epidermal regions after ATRA application was first documented in 1987; however, stem cell-level changes remained largely unexplored. This study demonstrates that ATRA treatment leads to reversible changes which decrease the fast-cycling epidermal compartment while expanding the slow-cycling one. ATRA biases both slow- and fast-cycling epidermal stem cell populations toward differentiation, with the remaining Slc1a3-CreER+ fast-cycling clones in the basal layers biasing to the slow-cycling lineage. Similar changes in slow- and fast-cycling epidermal stem cell populations are also evident in human primary cultures in vitro. These findings shed light on the role of retinoic acid signaling in regulating the balance of epidermal stem cell heterogeneity and lineages.

## INTRODUCTION

Retinoic acid (RA) and its derivatives have a variety of effects on skin and are widely used in the treatment of photoaging, intrinsic aging, and inflammatory diseases such as psoriasis (Li et al., 2017; Shao et al., 2017; W. Wang et al., 2020). All-trans retinoic acid (ATRA), a biologically active form of retinoid, is synthesized from dietary or topical retinol and retinyl esters (Szymański et al., 2020). RA levels are tightly controlled in tissues through biosynthesis and degradation. ATRA exerts its effects via the RA signaling pathway (Gur et al., 2022; Kurlandsky et al., 1996; Manolescu et al., 2010; Okano et al., 2012a).

The skin comprises the interfollicular epidermis and its appendages, including hair follicles, sebaceous glands, and sweat glands. RA signaling plays a crucial role in regulating epidermal cell proliferation and differentiation through RARγ/RXRα heterodimers and binding to the retinoic acid response element, partially mediated by the EGFR signaling pathway (Pasonen-Seppänen et al., 2008). RA signaling is crucial for epidermal maturation, skin barrier integrity, and hair growth, as demonstrated by mouse genetic models (Calléja et al., 2006; Okano et al., 2012b; Saitou et al., 1995), as well as pharmacological treatments of RAR/RXR antagonists or agonists (Calléja et al., 2006; Ishikawa et al., 2022). Recent studies have shed light on the cellular and molecular mechanisms of RA signaling in the regulation of hair follicle stem cells (HFSCs). The components necessary for RA production and degradation are precisely controlled throughout the hair cycle, ensuring the appropriate signaling activity of RA through the modulation of biological rhythms and Notch signaling pathways (Everts, 2012; Goggans et al., 2024). RA signaling facilitates HFSC plasticity by interacting with BMP and Wnt signaling, and regulates stem cell identity and lineage differentiation during homeostasis and injury repair (Tierney et al., 2024). RA signaling also plays a crucial role in enhancing HFSC non-professional phagocytotic ability (Stewart et al., 2024). Thus, ATRA serves as an essential modulator of cellular behavior, facilitating proper HFSC function and maintaining tissue integrity.

Mouse tail skin consists of anatomically distinct structures known as the interscale and scale, replenished by slowand fast-cycling epidermal stem cell populations differentiated by Dlx1 and Slc1a3 genetic markings, respectively (Sada et al., 2016). In homeostasis, slowand fast-cycling epidermal stem cells produce unique differentiation lineages: orthokeratotic K10+ lineage in the interscale, and parakeratotic K31/K36+ lineage in the scale (Fig. 1A). These stem cell populations alter their behavior guided by physiological and pathological cues, contributing to tissue remodeling. Skin injury induces inter-territorial migration of both slow- and fast-cycling epidermal stem cells (Sada et al., 2016), while skin aging results in the depletion of fast-cycling stem cells and a reduction in scale size (Raja et al., 2022). Schweizer et al. reported in 1987 that topical ATRA application causes a gradual reduction in scale size, with its conversion to interscale characterized by differential keratin expressions (Schweizer et al., 1987). However, the effects of RA signaling on slowand fast-cycling epidermal stem cell populations were not studied due to the lack of appropriate markers and lineage tracing tools. This study utilizes Dlx1 and Slc1a3 markers for in vivo lineage tracing, and addresses the effect of ATRA on the population balance and fate of epidermal stem cells.

**Figure 1.**
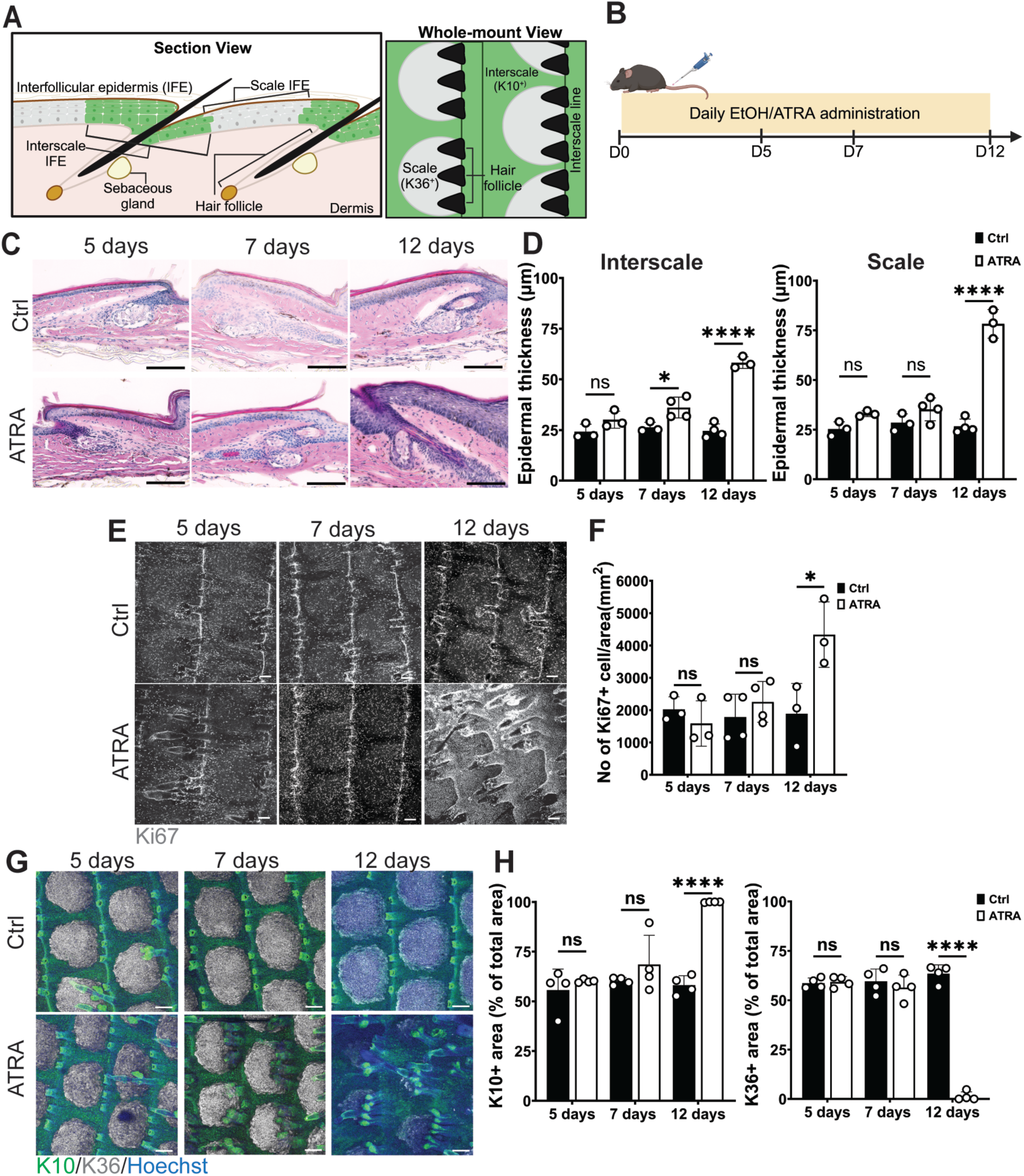
Impact of topical ATRA application on epidermal turnover and stem cell compartments in mouse tail skin. (**A**) Schematic representation of the interfollicular epidermis of mouse tail skin, shown in section and whole-mount views. Slow-cycling epidermal stem cells located in the interscale region (green). Fast-cycling stem cells located in the scale region (gray). The interscale line, where hair follicles penetrate, is abundant in Slc1a3-CreER-labeled cells. (**B**) Experimental scheme for ethanol (EtOH) (control: Ctrl) or ATRA topical application to mouse tail skin. Skin samples analyzed one day after final treatment. (**C**) Hematoxylin and eosin staining of skin sagittal sections. Scale bars: 150 µm. (**D**) Quantifications of epidermal thickness in interscale and scale regions. Data presented as mean ± S.D. Two-way ANOVA used for statistical analysis. *N* = 3, \**P* < 0.05, \*\*\*\**P* < 0.0001, ns; not significant. (**E, F**) Whole-mount staining of Ki67 (gray, a proliferation marker) and their quantifications. Scale bar: 100 µm. Two-way ANOVA used for statistical analysis. 5 and 12 days treatment; *N* = 3. 7 days treatment; *N* = 4, \**P* < 0.05, ns; not significant. (**G, H**) Whole-mount staining of K10 (green, interscale compartment) and K36 (gray, scale compartment) and quantifications. Blue, Hoechst (nucleus). Scale bars: 100 µm. Areas positive for K10 or K36 normalized to the total area, consisting of one interscale and one scale structure. Data presented as mean ± S.D. Two-way ANOVA used for statistical analysis. *N* = 3, \*\*\*\**P* < 0.0001, ns; not significant. Cartoons created with BioRender.com for panels (A, B).

## RESULTS

### ATRA treatment transiently skews the balance of slowand fast-cycling epidermal stem cell populations

To investigate the effect of ATRA on the balance of slow- and fast-cycling epidermal stem cell populations, 30 µg ATRA was topically applied to mouse tail skin (Fig. 1B). This led to epidermal hyperplasia, and converted parakeratotic to orthokeratotic scale regions in a dose-dependent manner; with repeated applications of the 30 µg dose, over 95% of skin areas showed this conversion (Schweizer et al., 1987). At 5 days, ATRA treatment did not cause any gross or histological changes in the skin (Fig. 1C, D. After 7 days, the interscale regions in ATRA-treated mice showed thicker epidermis than those in control mice (Fig. 1C, D). Over 12 days, epidermal thickness increased in both interscale and scale regions, with enhanced epidermal proliferation observed in ATRA-treated mice (Fig. 1C–F).

To examine the changes in epidermal stem cell compartments with ATRA treatment, whole-mount staining was performed to visualize slow-cycling (K10+ interscale) and fast-cycling (K36+ scale) lineages. At 5 and 7 days, K10+ interscale and K36+ scale areas were unaffected by ATRA. However, after 12 days, the K10+ interscale area significantly expanded, while the K36+ scale area decreased (Fig. 1G, H). These findings, consistent with those reported by Schweizer et al., suggest that ATRA disrupts the balance of slow- and fast-cycling epidermal stem cell populations in mouse skin, impacting the size of interscale and scale compartments.

To determine whether the observed changes in epidermal stem cells by ATRA were reversible, treatment was discontinued after 12 days, allowing the mice to recover (Fig. 2A). Epidermal thickness and scale compartments began to recover within 2 weeks of discontinuation of ATRA treatment, and scale compartments returned to their original size by Week 4 (Fig. 2B–E). These data suggest that the imbalance of scale/interscale regions due to ATRA treatment is not permanent; rather, the plasticity of epidermal stem cells is transiently affected when tissue RA levels are elevated.

**Figure 2.**
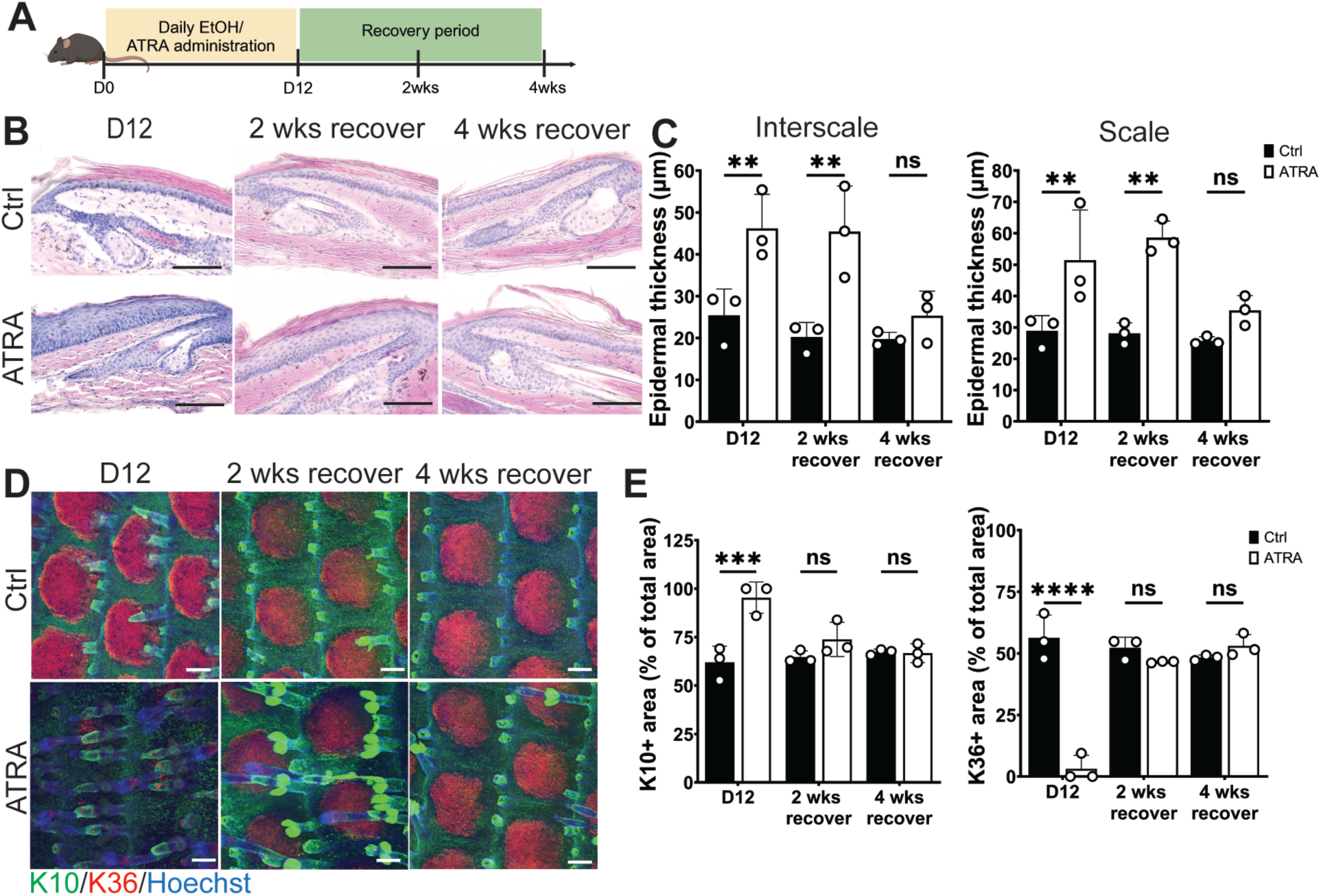
Recovery process after discontinuation of ATRA. (**A**) Experimental scheme. Mice treated with ethanol (EtOH) (control: Ctrl) or ATRA for 12 days, defined as day 12 (D12). Mice sacrificed at 2 and 4 weeks (wks) after treatment cessation. (**B**) Hematoxylin and eosin staining of skin sagittal sections. Scale bars: 150 µm. (**C**) Quantifications of epidermal thickness in interscale and scale regions. Data presented as mean ± S.D. Two-way ANOVA used for statistical analysis. *N* = 3, \*\**P* < 0.01, ns; not significant. (**D, E**) Whole-mount staining of K10 (green, interscale compartment) and K36 (gray, scale compartment) and quantifications. Blue, Hoechst (nucleus). Scale bars: 100 µm. Areas positive for K10 or K36 normalized to the total area, consisting of one interscale and one scale structure. Data presented as mean ± S.D. Two-way ANOVA used for statistical analysis. *N* = 3, \*\*\**P* < 0.001, ****P < 0.0001, ns; not significant. Cartoon created with BioRender.com for panel (A).

### ATRA alters the slowand fast-cycling epidermal stem cell markers in human primary keratinocytes

Next, to investigate whether the imbalance of epidermal stem cells induced by ATRA is cell-autonomous and also occurs in humans, human primary keratinocytes (epidermal stem/progenitor cells) were treated with ATRA. At 48 hours after ATRA treatment, there was a significant decrease in the basal marker *KRT14* and the early differentiation marker *KRT10* (Fig. 3B). In contrast, the late differentiation markers involucrin (*IVL*) and filaggrin (*FLG*) were elevated, suggesting that ATRA accelerates differentiation toward late stages (Fig. 3B).

**Figure 3.**
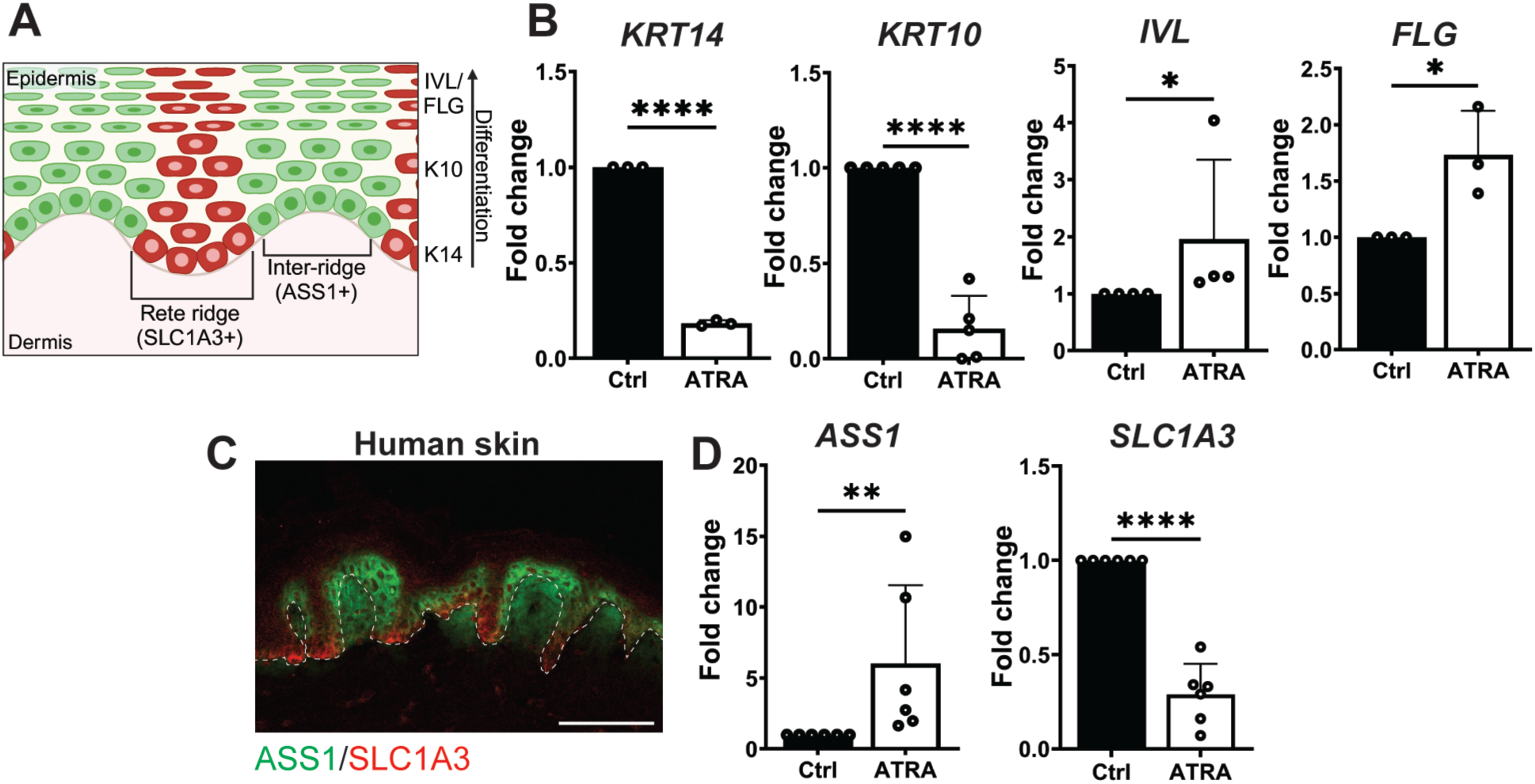
Alteration of epidermal stem cell markers in primary human keratinocytes. (**A**) Schematic representation of human skin. (**B**) Quantitative RT-PCR analysis of primary human keratinocytes treated with DMSO (control: Ctrl) or ATRA for 48 hours. Data presented as mean ± S.D. Two-tailed t-test used for statistical analysis. *KRT14*; *N* = 3. *KRT10*; *N* = 5. *IVL*; *N* = 4. *FLG*; *N* = 6, \**P* < 0.05, \*\*\**P* < 0.0001. (**C**) Section immunostaining of ASS1 (green, inter-ridge marker) and Slc1a3 (red, rete ridge marker) in human abdominal skin. White dashed line represents epidermal–dermal boundary. Scale bars: 100 µm. (D) Quantitative RT-PCR analysis of primary human keratinocytes treated with DMSO (control: Ctrl) or ATRA for 48 hours. Data presented as mean ± S.D. Two-tailed t-test used for statistical analysis. *N* = 6, \*\**P* < 0.005, \*\*\*\**P* < 0.0001. Cartoon created with BioRender.com for panel (A).

Recent single-cell RNA-seq analyses have shown similarities in the cell populations, lineages, and marker expressions of epidermal stem cells of mouse and human skin (Ghuwalewala et al., 2022). Notably, ASS1 and SLC1A3 exhibited heterogeneous expression patterns corresponding to the inter-ridge and rete ridge structures (Fig. 3A) (Ghuwalewala et al., 2022; Sada et al., 2016; S. Wang et al., 2020), as shown by immunostaining of human skin (Fig. 3C). This suggests an analogous relationship with the interscale and scale structures in mouse tail skin. Based on the staining patterns of these markers in vivo human skin, *ASS1* and *SLC1A3* were utilized as markers of slow- and fast-cycling epidermal stem cell populations, respectively, and the effects of ATRA on these markers in vitro were assessed. The presence of ATRA significantly altered the expression of these markers, increasing *ASS1* levels (indicative of slow-cycling epidermal stem cells) and suppressing *SLC1A3* expression (indicative of fast-cycling epidermal stem cells) (Fig. 3D). These results suggest that ATRA exerts a similar effect on human primary keratinocytes as on mice, biasing the balance of epidermal stem cells toward the slow-cycling population.

### A vitamin A deficient diet does not alter epidermal stem cell compartments

A vitamin A-deficient diet (VAD) was utilized to model systemic vitamin A deficiency in mice, and examine its effects on epidermal stem cell populations. Adult mice were fed a VAD for 11–16 weeks (Fig. 4A). The findings were consistent with previous reports (Cabezas-Wallscheid et al., 2017) demonstrating that when a VAD was introduced in adulthood, mouse body weight remained unchanged (Fig. 4B). However, white blood cell counts were reduced due to the vitamin A-dependent immune function and lymphoid tissue defects (Amimo et al., 2022; Mora et al., 2008) (Fig. 4C). There were no significant differences in skin histology (Fig. 4D, E) or interscale/scale size between regular and VAD-fed mice (Fig. 4F, G), suggesting that unlike HFSCs (Goggans et al., 2024) and hematopoietic stem cells (HSCs) (Cabezas-Wallscheid et al., 2017), blocking RA signaling locally or systemically has minimal impact on the heterogeneity of epidermal stem cells.

**Figure 4.**
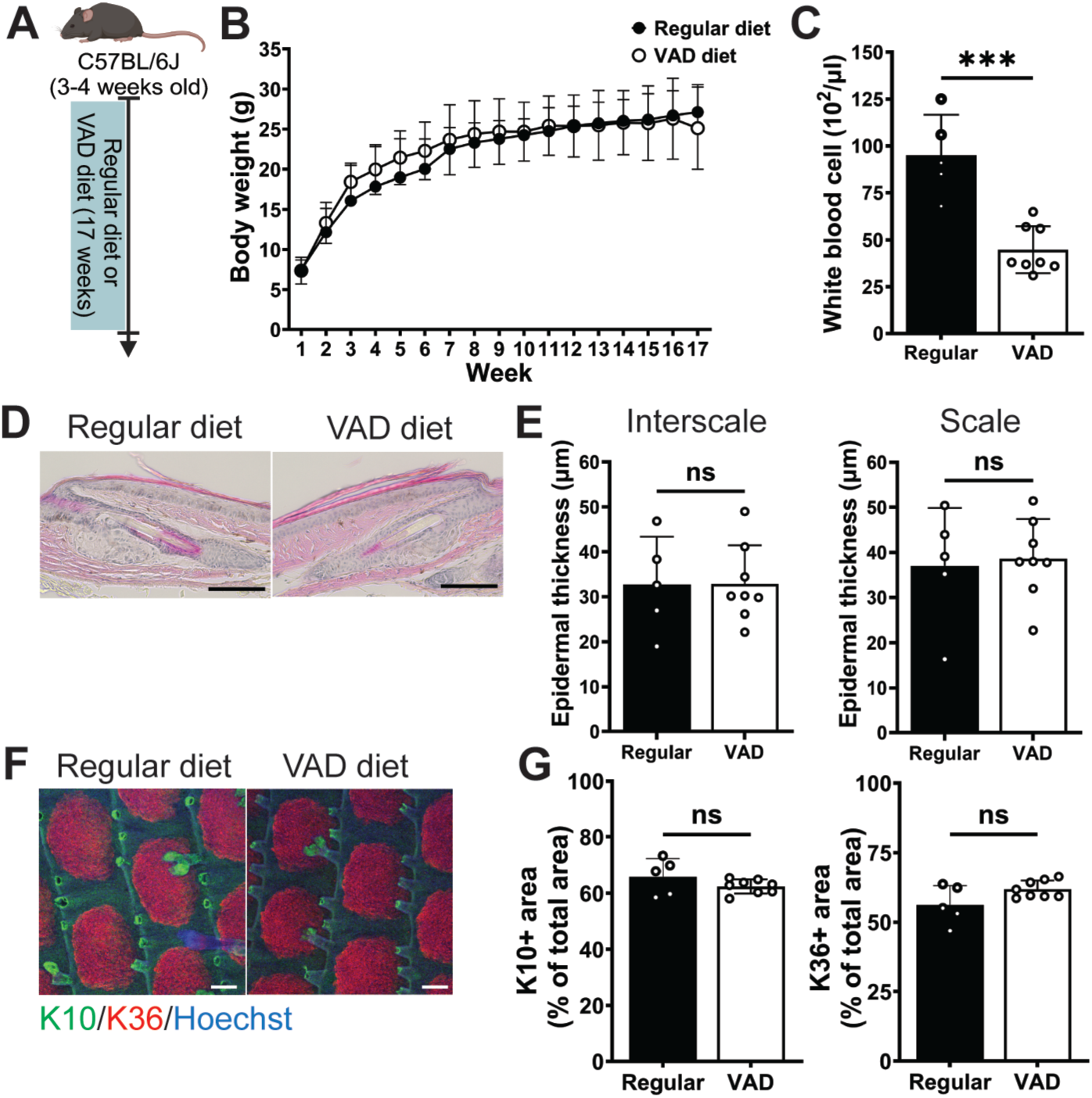
The effect of a vitamin A-deficient diet (VAD) on epidermal stem cell compartments. (**A**) Experimental scheme for administration of a regular diet or a VAD. (**B**) Body weight changes of regular diet-fed and VAD-fed mice. Data presented as mean ± S.D. Two-way ANOVA used for statistical analysis. No significant differences observed at any examined time point. Regular diet; *N* = 5. VAD; *N* = 8. (**C**) Number of white blood cells. Data presented as mean ± S.D. Two-tailed t-test used for statistical analysis. Regular diet; *N* = 5. VAD; *N* = 8. \*\*\**P* < 0.001. (**D**) Hematoxylin and eosin staining of skin sagittal sections. Scale bars: 150 µm. (**E**) Quantifications of epidermal thickness in interscale and scale regions. Data presented as mean ± S.D. Two-tailed t-test used for statistical analysis. Regular diet; *N* = 5. VAD; *N* = 8. ns; not significant. (**F, G**) Whole-mount staining of K10 (green, interscale compartment) and K36 (gray, scale compartment) and quantifications. Blue, Hoechst (nucleus). Scale bars: 100 µm. Areas positive for K10 or K36 normalized to the total area, consisting of one interscale and one scale structure. Data presented as mean ± S.D. Two-tailed t-test used for statistical analysis. Regular diet; *N* = 5. VAD; *N* = 8. ns; not significant. Cartoon created with BioRender.com for panel (A).

### ATRA promotes differentiation of both slowand fast-cycling epidermal stem cell populations

To investigate the effect of ATRA treatment on the size of interscale/scale compartments, Dlx1- and Slc1a3-CreER mice were used for lineage tracing (Fig. 5A) (Sada et al., 2016). At steady state, the interscale and scale regions are maintained by Dlx1+ slow-cycling epidermal stem cells and Slc1a3+ fast-cycling epidermal stem cells, respectively (Fig. 5B, left). There are two lineage behaviors of slow- and fast-cycling stem cell populations induced by ATRA which could explain the observed shrinkage of the scale region. One possibility is that the fast-cycling epidermal stem cell population is lost, allowing slow-cycling stem cells to dominate the entire epidermis (Fig. 5B, middle). Alternatively, the fast-cycling epidermal stem cell population may remain but alter fate, differentiating into the slow-cycling lineage (Fig. 5B, right).

**Figure 5.**
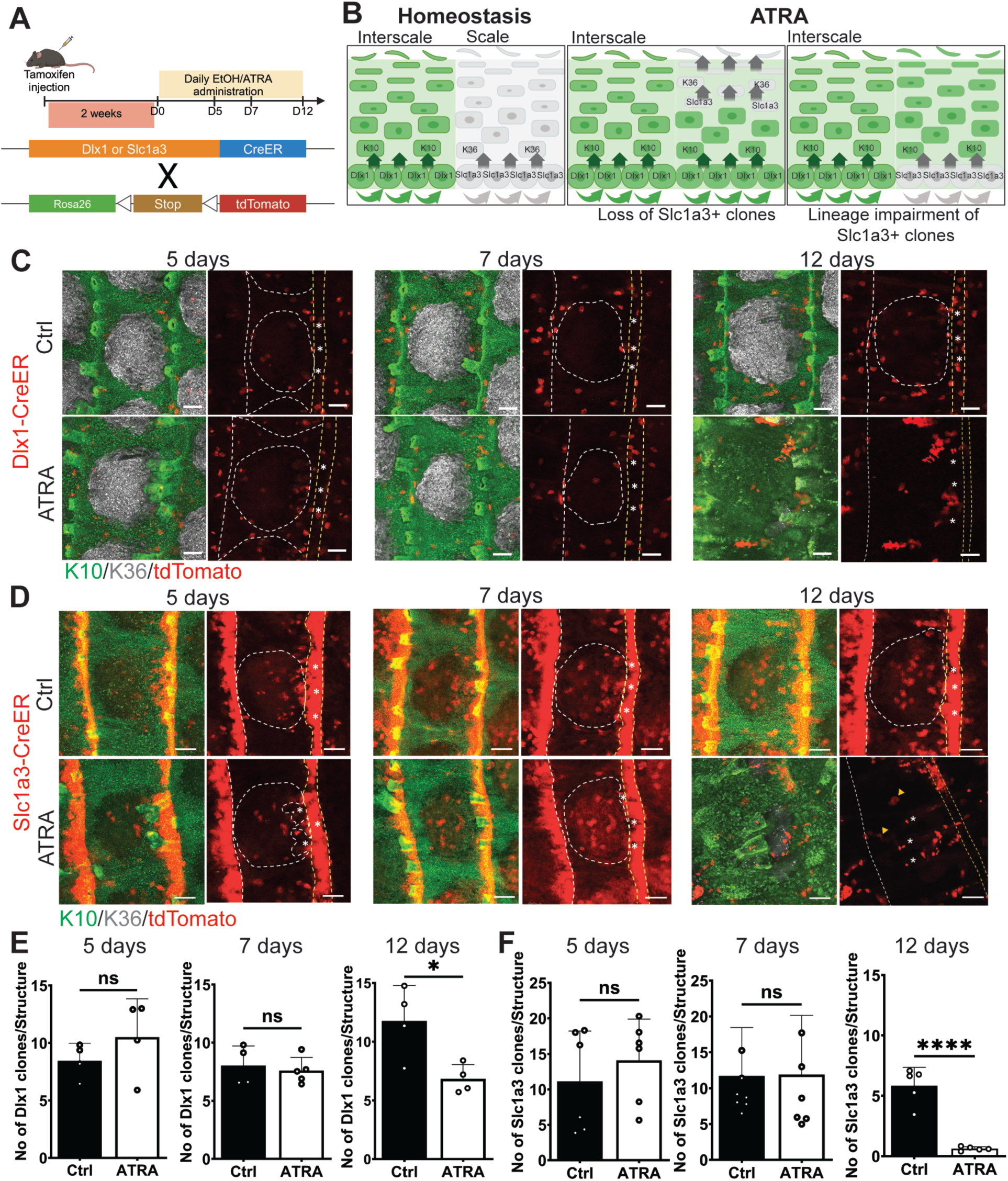
Behaviors of slowand fast-cycling epidermal stem cell clones with ATRA treatment. (**A**) Experimental scheme. Mice injected with a single dose of tamoxifen 2 weeks prior to treatment with ethanol (control: Ctrl) or ATRA. (**B**) Potential behaviors of Dlx1+ slow-cycling and Slc1a3+ fast-cycling epidermal stem cell populations under homeostatic or ATRA-treated conditions. (**C, D**) Whole-mount staining of Tomato (red), K10 (green, interscale compartment) and K36 (gray, scale compartment) and quantifications. Yellow dash lines indicate interscale line. White dash circles indicate scale compartment. Asterisks indicate hair follicles. Yellow arrowheads indicate Slc1a3+ clones overlap their expression with K10 in their original area. Scale bars: 100 µm. (**E**) Number of Dlx1-CreER+ clones per one interscale structure. Data presented as mean ± S.D. Two-tailed t-test used for statistical analysis. 5 days and 12 days treatment; *N* = 4. 7 days treatment; *N* = 5. \**P* < 0.05. ns; not significant. (**F**) Number of Slc1a3-CreER+ clones per one scale structure. Clones counted in original scale area, defined as the region beginning a 10-cell distance from the interscale region. Data presented as mean ± S.D. Two-tailed t-test used for statistical analysis. 5 days treatment; *N* = 6. 7 days treatment; *N* = 7. 12 days treatment; *N* = 5. *\*\*\*\*P* < 0.0001. ns; not significant. Cartoons created with BioRender.com for panels (A, B).

At days 5 and 7 of ATRA treatment (Fig. 5A), Dlx1+ clones were preferentially located in the interscale region, with no significant difference in clone numbers compared with controls (Fig. 5C, E). In contrast, Slc1a3+ clones were preferentially located in the scale and interscale line regions, as previously reported (Sada et al., 2016). The clone number was also unaffected. However, after 12 days, the number of Tomato+ clones in both Dlx1- and Slc1a3-CreER lineages was significantly decreased (Fig. 5C–F). To address the reason for this clone reduction in both lineages, the fate of slow- and fast-cycling stem cells with ATRA treatment on skin sections was examined. On days 5 and 7 of ATRA treatment, approximately 50% of Dlx1+ clones were found in the basal layer, similar to the control group. However, after 12 days, there was a significant increase in suprabasal (differentiating) clones, and a decrease in basal (self-renewing) clones to less than 25% (Fig. 6A, B). Similarly, Slc1a3+ clones were also primed for differentiation after 12 days of ATRA treatment (Fig. 6C, D). These results suggest that ATRA promotes differentiation of slowand fastcycling epidermal stem cell populations, leading to clone reduction in both lineages.

**Figure 6.**
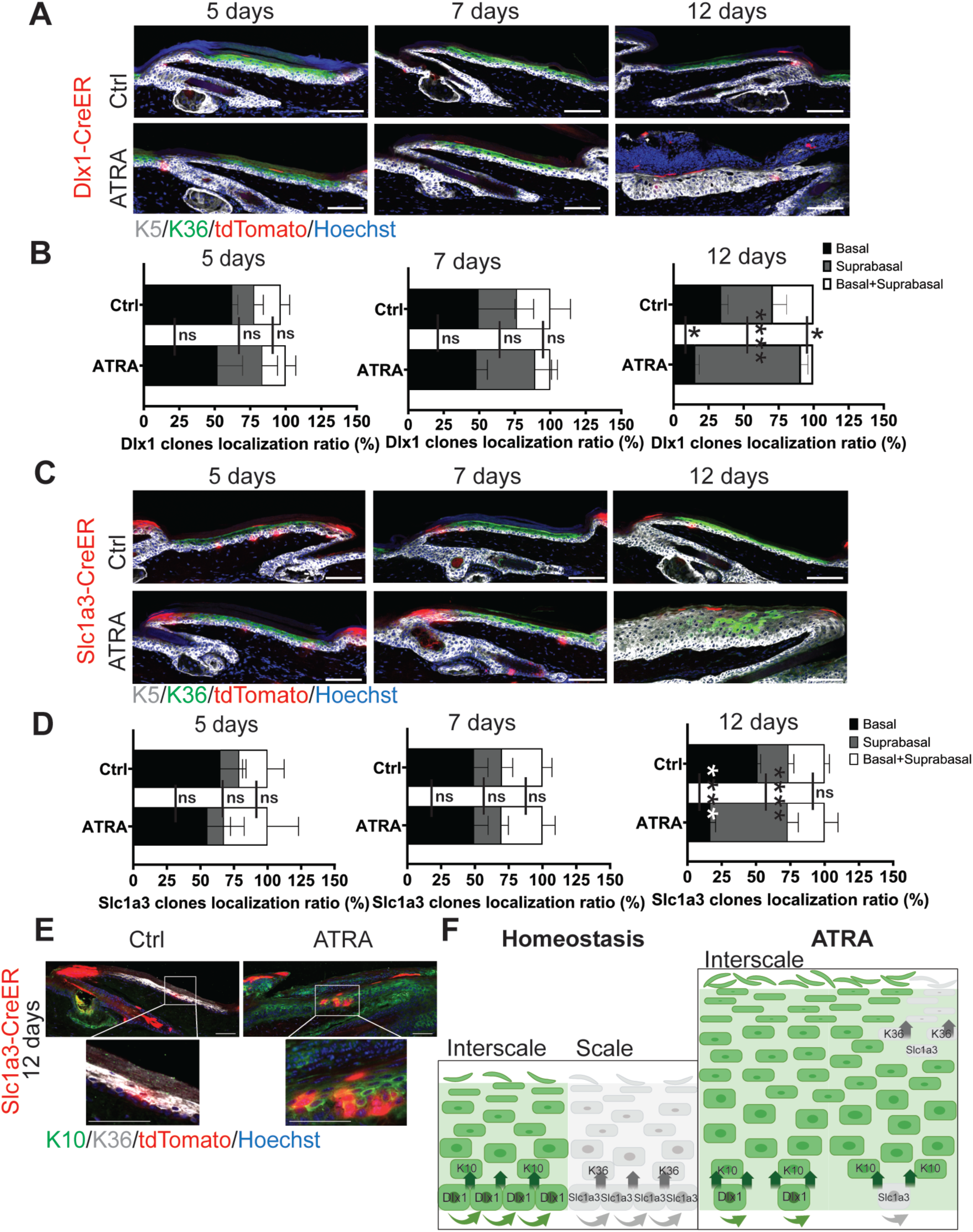
ATRA-induced changes in the fate of slowand fast-cycling epidermal stem cells. (**A**) Section immunostaining of Dlx1-CreER mice with K5 (gray, epidermal basal layer), K36 (gray, scale compartment), and Tomato (red). Blue, Hoechst (nucleus). Scale bars: 100 µm. (**B**) Percentage of Dlx1-CreER+ clones located in the (1) basal layer only (self-renewing clones, black bars); (2) suprabasal layer only (differentiating clones, gray bars); or (3) both basal and suprabasal layers (white bars). Quantification performed on Dlx1-CreER+ clones in the interscale region. Data presented as mean ± S.D. Two-way ANOVA used for statistical analysis. *N* = 3, \**P* < 0.05, \*\*\*\**P* < 0.0001. ns; not significant. (**C**) Section immunostaining of Slc1a3-CreER mice with K5, K36, Hoechst, and Tomato (red). Scale bar: 100 µm. (**D**) Percentage of Slc1a3-CreER+ clones located in the (1) basal layer only (self-renewing clones, black bars); (2) suprabasal layer only (differentiating clones, gray bars); or (3) both basal and suprabasal layers (white bars). Quantification performed on Slc1a3-CreER + clones in the scale region. Clones counted in original scale area, defined as the region beginning a 10-cell distance from the interscale region. Data presented as mean ± S.D. Two-way ANOVA used for statistical analysis. *N* = 3, \*\*\*\**P* < 0.0001. ns; not significant. (**E**) Section immunostaining of K10 (green, interscale compartment), K36 (gray, scale compartment), and Tomato (red). Blue, Hoechst (nucleus). Scale bars: 100 µm. (**F**) Schematic summarizing the results of the lineage tracing analysis. In the control condition, Dlx1-CreER+ slowcycling epidermal stem cells produce a K10+ interscale differentiation lineage (green); Slc1a3 produces a K36+ scale differentiation lineage (gray). With ATRA treatment, the entire epidermis becomes K10-positive. Dlx1- and Slc1a3-positive epidermal stem cell populations show a decrease in clones and a shift toward differentiation. Ectopic K10-positive cells appear in the Slc1a3 lineage, indicating conversion of the differentiation lineage into interscale. Cartoon created with BioRender.com for panel (F).

### ATRA biases fast-cycling epidermal stem cells toward slow-cycling lineage differentiation

To address the reason for the overall balance shift toward the slow-cycling lineage, the possibility of fate changes was addressed in both populations. After 12 days of ATRA treatment, when the K36+ scale region was depleted, Dlx1+ clones continued to express the K10 interscale marker, and remained at their original position without active migration toward the scale region (Fig. 5C). At this time point, some Slc1a3+ clones were observed at the center of the original scale position, which was originally positive for K36 but began expressing K10 (Fig. 5D), indicating lineage changes.

On skin sections, Slc1a3+ fast-cycling epidermal clones were found to gradually diminish through differentiation following ATRA treatment, leaving only a few in the basal layer (Fig. 6D). At 12 days of ATRA treatment, Slc1a3+ clones were found in both the basal and suprabasal layers within the original scale region by their position, indicating their self-renewal and differentiation abilities. Notably, these remaining Slc1a3+ clones co-expressed the K10 (interscale) marker in the upper layer, which was rarely observed in the control group (Fig. 6E). Altogether, these results suggest that ATRA skews both slow- and fast-cycling populations toward differentiation, with remaining fast-cycling epidermal basal cells biasing toward the K10+ slow-cycling lineage, causing the loss of scale structure in the interfollicular epidermis (Fig. 6F).

## DISCUSSION

RA and its derivatives have been widely used in medical and cosmetic skin formulations. Despite this clinical importance, the cellular mechanisms of ATRA action in the interfollicular epidermis— particularly its effects on heterogeneous stem cell populations—remain poorly understood. The current study revealed that ATRA impacts the balance of slow- and fast-cycling stem cell populations in the interfollicular epidermis of mouse skin and human keratinocytes. Lineage tracing analysis showed that ATRA not only promotes differentiation of epidermal stem cells, but also shifts fast-cycling populations toward slow-cycling lineages. Upon discontinuation of ATRA treatment, the skin returned to its original state, suggesting that the observed stem cell fate switching is temporary. Similar changes have been reported in HSCs, where ATRA facilitates the re-entry of active HSCs into quiescence, and the absence of ATRA hinders HSCs from recovering to a quiescent state (Cabezas-Wallscheid et al., 2017; Zhang et al., 2022). ATRA also regulates the plasticity and lineage of HFSCs (Tierney et al., 2024). These findings suggest that RA signaling plays a fundamental role in regulating stem cell heterogeneity and fate across different tissues.

This study found that both Dlx1+ and Slc1a3+ epidermal stem cell populations responded to ATRA and were biased toward differentiation. Furthermore, the scale structure which disappeared following ATRA administration, was restored upon discontinuation. These results suggest two possibilities: (1) the Slc1a3+ epidermal stem cell population may have transiently shifted to the interscale lineage, or (2) the newly formed scale structures may have originated from slow-cycling stem cells. The lineage analysis revealed that with ATRA treatment, some Slc1a3+ clones localized to the original scale position and expressed the K10 marker, suggesting that ATRA may induce a differentiation pathway from scale-to interscale-type. This transient plasticity of fast-cycling stem cells to slow-cycling lineages is also observed in mouse tail skin scratch wounds (Sada et al., 2016). The underlying molecular mechanism is not yet understood, but it is possible that Wnt and EGFR signaling, which promote the induction of scale lineage (Gomez et al., 2013), are suppressed by elevated RA signaling. Future studies should explore the process of stem cell fate switching in response to ATRA by combining cell lineage with single-cell level transcriptome/epigenome analysis over time.

In previous studies using the vitamin A deficiency model, pregnant female mice were fed a VAD which affected fetal embryonic development, leading to impaired barrier function, metaplasia, and altered hair formation (Girard et al., 2006; Huang et al., 1986; Klein-Szanto et al., 1982; Surman et al., 2020). In contrast, this study applied the conditions of a previous HSC investigation, administering the VAD postnatally to avoid embryonic effects. Under these conditions, while there were significant impacts on the functionality of HSCs (Cabezas-Wallscheid et al., 2017), no noticeable changes in interscale/scale size or skin histology were observed (Fig. 4). This suggests that endogenous RA may play a minimal role in regulating epidermal stem cell balance in adulthood, once stem cell populations are established. Further research is necessary to explore the potential effects of vitamin A deficiency on epidermal stem cell populations and skin regeneration under stress conditions.

While ATRA offers therapeutic benefits, high concentrations could lead to notable adverse effects (Lee et al., 2010; Szymański et al., 2020). In humans, topical RA treatment results in epidermal thickening due to increased epidermal proliferation (Kang et al., 1995), similar to findings of the current mouse study (Fig. 1C–F). However, prolonged treatment may eventually cause epidermal thinning due to the depletion and accelerated differentiation of epidermal stem cells. Additionally, this study found that ATRA skews the fates of slow- and fast-cycling epidermal stem cell populations, which mechanism had as yet not been elucidated. Future investigations should explore the potential adverse effects of elevated RA levels in the skin, and aim to develop targeted interventions minimizing side effects.

## MATERIALS AND METHODS

### Mice

All mouse experiments were conducted according to the guidelines for animal experimentation by the Institutional Animal Experiment Committee at Kumamoto University and Kyushu University. Experimental mice were housed in specific-pathogen-free environments at the Center for Animal Resources and Development, Kumamoto University, and the Laboratory of Embryonic and Genetic Engineering, Kyushu University. Wild-type (C57BL/6J) mice were purchased from Charles River Laboratories or Japan SLC, Inc. For lineage tracing experiments, Dlx1-CreER (C57BL/6J) (The Jackson Laboratory, no. 014551) (Taniguchi et al., 2011) or Slc1a-3CreER (C57BL/6J) (The Jackson Laboratory, no. 012586) mice were crossed with Rosa-tdTomato reporter mice (C57BL/6J) (The Jackson Laboratory, no. 007905) to obtain double transgenic mice. Both male and female mice were used in all experiments.

### ATRA treatment

ATRA (FUJIFILM Wako Pure Chemical Corporation) was dissolved in ethanol to a concentration of 30 µg per 100 µl ethanol. Two-month-old mice were treated with either 100 µl ethanol or 30 µg/100 µl ATRA, applied topically to tail skin daily for durations of 5, 7, and 12 days. Mice were sacrificed one day after the final treatment.

### VAD mice

C57BL/6J mice, aged 3–4 weeks, were fed either a regular diet or a VAD (Oriental Kobo Inc.) for durations of 17 weeks as previously described (Cabezas-Wallscheid et al., 2017). The mice were weighed weekly. Peripheral blood was collected via the retroorbital sinus using a heparin-coated capillary tube and transferred to EDTA-coated tubes. The white blood cells were measured with a Celltac-α according to the manufacturer’s instructions (MEK-6550, Nihon Kohden).

### Tamoxifen administration

Tamoxifen (Sigma T5648) was dissolved in corn oil. For the lineage tracing, Slc1a3-CreER/RosatdTomato and Dlx1-CreER/Rosa-tdTomato mice at 2 months of age were injected intraperitoneally with a single dose of tamoxifen at 100 µg/g and 50 µg/g of body weight, respectively.

### Hematoxylin and eosin staining

Tail skin samples were embedded in optimal cutting temperature (O.C.T.) compound (Sakura Finetek). Sections were cut to 10 µm thickness, dried at room temperature (RT) for 1 hour, and fixed with 4% paraformaldehyde (PFA) for 10 minutes. The sections were then stained with hematoxylin for 20 minutes and eosin for 5 minutes. After staining, the sections were dehydrated and mounted with Entellan new mounting medium (Sigma-Aldrich). Images were captured using an EVOS M5000 Imaging System (ThermoFisher Scientific). Epidermal thickness was quantified for at least six structures, each consisting of one scale and one interscale structure, using ImageJ software (NIH).

### Whole-mount immunostaining

Mouse tail skin was cut into 5 mm × 5 mm pieces and incubated in 20 mM EDTA/PBS on a shaker at 37°C for 2 hours. The epidermis was separated from the dermis and fixed with 4% PFA overnight at 4°C. Epidermal sheets were incubated in a blocking buffer (1% BSA, 2.5% donkey serum, 2.5% goat serum, 0.8% Triton in PBS) for 3 hours at RT on a shaker, then incubated with primary antibodies in blocking buffer overnight at RT. The following primary antibodies were used: mouse anti-K10 (1:500, Abcam no. ab9026); rabbit anti-K36 (1:300, Proteintech 14309-1-AP); rabbit anti-K14 (1:1,000, BioLegend no. 905304); and rat anti-Ki67 (1:100, Invitrogen no.14-5698-82). Mouse on Mouse (M.O.M.) kit reagents (Vector Laboratories, Inc.) were added when using a primary mouse antibody. Samples were washed four times for 1 hour with 0.2% Tween 20/PBS at RT, followed by incubation with a secondary antibody in blocking buffer overnight at 4°C. All secondary primary antibodies (Alexa 488, 555, and 647, Invitrogen) were diluted at 1:500 to 1:300. After washing, samples were stained with Hoechst (Sigma-Aldrich) for 1 hour at RT and mounted.

### Immunostaining of skin sections

Mouse tail skin sections were fixed with 4% PFA for 10 minutes at RT and incubated with blocking solution (1% bovine serum albumin, 2.5% normal goat serum, 2.5% normal donkey serum, 2% gelatin, and 0.1% Triton X-100 in PBS) for 1 hour at RT. The sections were incubated with primary antibodies overnight at 4°C. After washing, sections were incubated with secondary antibodies for 1 hour at RT. The sections were washed and stained with Hoechst for 10 minutes at RT and mounted. The following primary antibodies were used: rabbit anti-K14 (1:500, BioLegend, 905301); mouse anti-K10 (1:500, Abcam, ab9026); rabbit anti-K36 (1:300, Proteintech, 14309-1-AP); and chicken anti-K5 (1:500, BioLegend, 905904). All secondary primary antibodies (Alexa 488, 555, and 647, Invitrogen) were diluted at 1:300. M.O.M. kit reagents (Vector Laboratories) were added when using a primary mouse antibody.

### Human skin samples

Human abdominal skin was purchased from CTI-Biotech (Lyon, France) (SK534, SK545, SK548, and SK550). These samples were collected from anonymous donor patients aged 20–40 years, with informed consent. The consent and collection procedures were in accordance with European standards and applicable ethical guidelines. Skin sections were cut to 10 µm thickness and stained as described above. The primary antibodies used were as follows: rabbit anti-ASS1 (1:500, Cell Signaling, 70720S); guinea pig anti-Slc1a3 (1:100, Frontier Institute, GLAST-GP-Af1000); and chicken anti-K5 (1:500, BioLegend, 905904). Secondary antibodies (Alexa 488, 555, and 647, Invitrogen) were diluted at 1:300.

### Imaging and quantification

All whole-mount or section staining samples were observed using a confocal microscope (Nikon A1 HD25 or Zeiss LSM900), with images captured and analyzed through NIS Elements Imaging or Zen 3 (Blue Edition) software. Images were adjusted using Adobe Photoshop 2023. K10 and K36 positive areas, the number of tdTomato clones, and Ki67 positive cells in whole-mount samples were measured and counted from 6 to 12 images using ImageJ software (NIH). The number of tdTomato clones from mouse sections was manually counted from 20 to 30 structures. Statistical analyses and graphs were generated using GraphPad Prism 9 software.

### Primary keratinocyte culture

Neonatal human primary keratinocytes (KER112002, KER112005, and KER112006, BIOPREDIC International) were cultured in CnT-PR medium (CELLnTEC) on collagen IV (Sigma-Aldrich) pre-coated 100 mm dishes until they reached 70–80% confluency. Cells were passaged using 2 ml Accutase (ICT) and 1 ml Accumax (ICT) per 100 mm dish for 15–20 minutes at RT, then seeded onto collagen IV-coated 12-well plates at a density of 50,000 cells per well. Cells were cultured in a CnT-PR medium at 37°C in 5% CO2 for 24 hours. The medium was then replaced with a CnT-PR medium with or without 5 µM ATRA, and cells cultured for 48 hours. Cells were subsequently collected for further experiments.

### Quantitative RT-PCR

Total RNA from cultured human keratinocytes was isolated utilizing the RNeasy Micro Kit (QIAGEN) in accordance with the manufacturer’s protocol, and cDNA synthesis was performed with iScript Reverse Transcription Supermix (Bio-Rad). Quantitative RT-PCR was conducted on a LightCycler 96 System (Roche) using the FastStart Essential DNA Green Master (Roche) with 5 ng of cDNA per well. Target gene expression was normalized to the ®-actin housekeeping gene. The primer sequences are as follows: *SLC1A3* forward (F) 5’-AGCAGGGAGTCCGTAAACGC-3’ and reverse (R) 5’-TGGTCGGAGGGTAAATCCAAGG-3’; *ASS1* F 5’-TCTACAAC-CGGTTCAAGGGC-3’ and R 5’-TCCAGGATTCCAGCCTCGTA-3’, *KRT14* F 5’-AGATGATTGGCAGCGTGGAG-3’ and R 5’-AACTGGGAGGAGGAGAGGTG-3’; *KRT10* F 5’-TCGGGCTCTGGAAGAATCAA-3’ and R 5’-CTGAAGCAGGATGTTGGCATT-3’; *IVL* F 5’-GAACAGCAGGAAAAGCACCT-3’ and R 5’-CTGGTTGAATGTCTTGGACCT-3’; *FLG* F 5’-CAGTGAGGCATACCCAGAGG-3’ and R 5’-ACTGGCTGTATCGCGGTGAG-3’; *ACTB* F 5’-AGAGATGGCCACGGCTGCTT-3’ and R 5’-ATTTGCGGTGGACGATGGAG-3’.

## Acknowledgments

We thank the Center for Animal Resources and Development (CARD) at Kumamoto University, the Laboratory of Embryonic and Genetic Engineering at Kyushu University, and T. Keida (Kumamoto University) and N. Imai (Kyushu University) for mouse care. We also thank the International Core-facility of Advanced Life Science at Kumamoto University and the Research Promotion Unit at Kyushu University for their facility support. We thank G. Sashida (Kumamoto University) for technical assistance with blood cell measurement. We also thank Dr. A. Yesbolatova (Kyushu University) for critical reading of this manuscript. This work was supported by Grant-in-Aid for Scientific Research (B) (24K02035, 20H03266) (to A.S.), Grant-in-Aid for Challenging Research (Exploratory) (24K21973) (to A.S.), and the Takeda Science Foundation (to A.S.). We also acknowledge support from the RMUTT Human Resource Development Scholarship (to T.D.).

## CRediT Statement (Author Contributions)

Conceptualization: A.S.; Investigation: T.D. and M.I.; Formal Analysis: T.D.; Validation: T.D., A.S., and N.C.-W.; Resources: A.S.; Methodology: A.S. and N.C.-W.; Supervision: A.S.; Funding Acquisition: A.S.; Visualization: T.D.; Writing—original draft preparation: T.D.; Writing—review and editing: A.S., N.C.-W., and M.I.

## Disclosure and Competing Interests Statement

The authors declare no conflicts of interest.

## Declaration of Generative AI and AI-Assisted Technologies in the Writing Process

During the preparation of this manuscript, the authors used DeepL, Grammarly, and ChatGPT to assist with language clarity and readability. Following the use of these tools, the authors thoroughly reviewed and edited the content to ensure accuracy. The authors take full responsibility for the final version of the published article.

## Notes

### Competing Interest Statement

The authors have declared no competing interest.

## References

Amimo JO, Michael H, Chepngeno J, Raev SA, Saif LJ, Vlasova AN. Immune Impairment Associated with Vitamin A Deficiency: Insights from Clinical Studies and Animal Model Research. Nutrients 2022;14:5038. 10.3390/nu14235038.

Cabezas-Wallscheid N, Buettner F, Sommerkamp P, Klimmeck D, Ladel L, Thalheimer FB, et al. Vitamin A-Retinoic Acid Signaling Regulates Hematopoietic Stem Cell Dormancy. Cell 2017;169:807–823.e19. 10.1016/j.cell.2017.04.018.

Calléja C, Messaddeq N, Chapellier B, Yang H, Krezel W, Li M, et al. Genetic and pharmacological evidence that a retinoic acid cannot be the RXR-activating ligand in mouse epidermis keratinocytes. Genes Dev 2006;20:1525–38. 10.1101/gad.368706.

Everts HB. Endogenous retinoids in the hair follicle and sebaceous gland. Biochim Biophys Acta 2012;1821:222–9. 10.1016/j.bbalip.2011.08.017.

Ghuwalewala S, Lee SA, Jiang K, Baidya J, Chovatiya G, Kaur P, et al. Binary organization of epidermal basal domains highlights robustness to environmental exposure. EMBO J 2022;41:e110488. 10.15252/embj.2021110488.

Girard C, Dereure O, Blatière V, Guillot B, Bessis D. Vitamin A Deficiency Phrynoderma Associated with Chronic Giardiasis. Pediatr Dermatol 2006;23:346–9. 10.1111/j.1525-1470.2006.00261.x.

Goggans KR, Belyaeva OV, Klyuyeva AV, Studdard J, Slay A, Newman RB, et al. Epidermal retinol dehydrogenases cyclically regulate stem cell markers and clock genes and influence hair composition. Commun Biol 2024;7:1–12. 10.1038/s42003-024-06160-2.

Gomez C, Chua W, Miremadi A, Quist S, Headon DJ, Watt FM. The Interfollicular Epidermis of Adult Mouse Tail Comprises Two Distinct Cell Lineages that Are Differentially Regulated by Wnt, Edaradd, and Lrig1. Stem Cell Rep 2013;1:19–27. 10.1016/j.stemcr.2013.04.001.

Gur M, Bendelac-Kapon L, Shabtai Y, Pillemer G, Fainsod A. Reduced Retinoic Acid Signaling During Gastrulation Induces Developmental Microcephaly. Front Cell Dev Biol 2022;10. 10.3389/fcell.2022.844619.

Huang FL, Roop DR, De Luca LM. Vitamin A deficiency and keratin biosynthesis in cultured hamster trachea. Vitro Cell Dev Biol J Tissue Cult Assoc 1986;22:223–30. 10.1007/BF02623307.

Ishikawa S, Nikaido M, Otani T, Ogata K, Iida H, Inai Y, et al. Inhibition of retinoid X receptor improved the morphology, localization of desmosomal proteins and paracellular permeability in three-dimensional cultures of mouse keratinocytes. Microsc Oxf Engl 2022;71:152–60. 10.1093/jmicro/dfac007.

Kang S, Duell EA, Fisher GJ, Datta SC, Wang Z-Q, Reddy AP, et al. Application of Retinol to Human Skin In Vivo Induces Epidermal Hyperplasia and Cellular Retinoid Binding Proteins Characteristic of Retinoic Acid but Without Measurable Retinoic Acid Levels or Irritation. J Invest Dermatol 1995;105:549–56. 10.1111/1523-1747.ep12323445.

Klein-Szanto AJP, Martin D, Sega M. Hyperkeratinization and hyperplasia of the forestomach epithelium in vitamin A deficient rats. Virchows Arch B 1982;40:387–94. 10.1007/BF02932880.

Kurlandsky SB, Duell EA, Kang S, Voorhees JJ, Fisher GJ. Auto-regulation of Retinoic Acid Biosynthesis through Regulation of Retinol Esterification in Human Keratinocytes*. J Biol Chem 1996;271:15346–52. 10.1074/jbc.271.26.15346.

Lee JE, Chang JY, Lee SE, Kim MY, Lee JS, Lee MG, et al. Epidermal Hyperplasia and Elevated HB-EGF are More Prominent in Retinoid Dermatitis Compared with Irritant Contact Dermatitis Induced by Benzalkonium Chloride. Ann Dermatol 2010;22:290–9. 10.5021/ad.2010.22.3.290.

Li Z, Niu X, Xiao S, Ma H. Retinoic acid ameliorates photoaged skin through RAR-mediated pathway in mice. Mol Med Rep 2017;16:6240–7. 10.3892/mmr.2017.7336.

Manolescu DC, El-Kares R, Lakhal-Chaieb L, Montpetit A, Bhat PV, Goodyer P. Newborn Serum Retinoic Acid Level Is Associated With Variants of Genes in the Retinol Metabolism Pathway. Pediatr Res 2010;67:598–602. 10.1203/PDR.0b013e3181dcf18a.

Mora JR, Iwata M, von Andrian UH. Vitamin effects on the immune system: vitamins A and D take centre stage. Nat Rev Immunol 2008;8:685–98. 10.1038/nri2378.

Okano J, Lichti U, Mamiya S, Aronova M, Zhang G, Yuspa SH, et al. Increased retinoic acid levels through ablation of Cyp26b1 determine the processes of embryonic skin barrier formation and peridermal development. J Cell Sci 2012a;125:1827–36. 10.1242/jcs.101550.

Okano J, Levy C, Lichti U, Sun H-W, Yuspa SH, Sakai Y, et al. Cutaneous Retinoic Acid Levels Determine Hair Follicle Development and Downgrowth. J Biol Chem 2012b;287:39304–15. 10.1074/jbc.M112.397273.

Pasonen-Seppänen SM, Maytin EV, Törrönen KJ, Hyttinen JMT, Hascall VC, MacCallum DK, et al. All-trans retinoic acid-induced hyaluronan production and hyperplasia are partly mediated by EGFR signaling in epidermal keratinocytes. J Invest Dermatol 2008;128:797–807. 10.1038/sj.jid.5701098.

Raja E, Changarathil G, Oinam L, Tsunezumi J, Ngo YX, Ishii R, et al. The extracellular matrix fibulin 7 maintains epidermal stem cell heterogeneity during skin aging. EMBO Rep 2022;23:e55478. 10.15252/embr.202255478.

Sada A, Jacob F, Leung E, Wang S, White BS, Shalloway D, et al. Defining the cellular lineage hierarchy in the interfollicular epidermis of adult skin. Nat Cell Biol 2016;18:619–31. 10.1038/ncb3359.

Saitou M, Sugai S, Tanaka T, Shimouchi K, Fuchs E, Narumiya S, et al. Inhibition of skin development by targeted expression of a dominant-negative retinoic acid receptor. Nature 1995;374:159–62. 10.1038/374159a0.

Schweizer J, Fürstenberger G, Winter H. Selective suppression of two postnatally acquired 70 kD and 65 kD keratin proteins during continuous treatment of adult mouse tail epidermis with vitamin A. J Invest Dermatol 1987;89:125–31. 10.1111/1523-1747.ep12470544.

Shao Y, He T, Fisher GJ, Voorhees JJ, Quan T. Molecular basis of retinol anti-ageing properties in naturally aged human skin in vivo. Int J Cosmet Sci 2017;39:56–65. 10.1111/ics.12348.

Stewart KS, Abdusselamoglu MD, Tierney MT, Gola A, Hur YH, Gonzales KAU, et al. Stem cells tightly regulate dead cell clearance to maintain tissue fitness. Nature 2024;633:407–16. 10.1038/s41586-024-07855-6.

Surman SL, Penkert RR, Sealy RE, Jones BG, Marion TN, Vogel P, et al. Consequences of Vitamin A Deficiency: Immunoglobulin Dysregulation, Squamous Cell Metaplasia, Infectious Disease, and Death. Int J Mol Sci 2020;21:5570. 10.3390/ijms21155570.

Szymański Ł, Skopek R, Palusińska M, Schenk T, Stengel S, Lewicki S, et al. Retinoic Acid and Its Derivatives in Skin. Cells 2020;9:2660. 10.3390/cells9122660.

Taniguchi H, He M, Wu P, Kim S, Paik R, Sugino K, et al. A resource of Cre driver lines for genetic targeting of GABAergic neurons in cerebral cortex. Neuron 2011;71:995–1013. 10.1016/j.neuron.2011.07.026.

Tierney MT, Polak L, Yang Y, Abdusselamoglu MD, Baek I, Stewart KS, et al. Vitamin A resolves lineage plasticity to orchestrate stem cell lineage choices. Science 2024;383:eadi7342. 10.1126/science.adi7342.

Wang S, Drummond ML, Guerrero-Juarez CF, Tarapore E, MacLean AL, Stabell AR, et al. Single cell transcriptomics of human epidermis identifies basal stem cell transition states. Nat Commun 2020;11:4239. 10.1038/s41467-020-18075-7.

Wang W, Shu G, Lu K, Xu X, Sun M, Qi J, et al. Flexible liposomal gel dual-loaded with alltrans retinoic acid and betamethasone for enhanced therapeutic efficiency of psoriasis. J Nanobiotechnology 2020;18:80. 10.1186/s12951-020-00635-0.

Zhang YW, Mess J, Aizarani N, Mishra P, Johnson C, Romero-Mulero MC, et al. Hyaluronic acid–GPRC5C signalling promotes dormancy in haematopoietic stem cells. Nat Cell Biol 2022;24:1038–48. 10.1038/s41556-022-00931-x.

